# A purification-free nucleic acid amplification platform for diverse samples

**DOI:** 10.1101/2025.05.02.651961

**Authors:** Jay Bhakti Kapadia, Kadidia Dite Selly N’Diaye, Jamal Daoud, Jonathan Perreault

## Abstract

An enzyme-free nucleic acid amplification method based on toehold-mediated strand displacement reaction (TMSDR) was evaluated under a variety of conditions with the aim of eliminating conventional purification steps and streamlining diagnostic workflows. By operating directly in lysis buffers, the TMSDR assay enhances target recovery and confers protection against nuclease degradation. Amplification performance was examined in the presence of diverse denaturing chemicals, lysis buffers, and sample matrices—including blood, saliva, wastewater, and soil—and the results demonstrated broad versatility and robust amplification under most conditions. The assay-maintained efficacy even in the presence of common PCR inhibitors, such as polyphenols in plant extracts, immunoglobulin G in blood, and complex constituents in environmental samples. Furthermore, a proof-of-concept assay targeting the 16S rRNA of *Escherichia coli* DH5α was established using specifically designed probe and displacer sequences, with specificity confirmed by the absence of amplification in mutated target controls. Collectively, these findings underscore the potential of the TMSDR assay as a highly adaptable and efficient alternative for rapid nucleic acid detection in point-of-care and field applications. This enzyme-free, purification-independent platform opens the door for rapid diagnostic development in resource-limited settings

## Introduction

Amplification techniques are essential for detecting and quantifying nucleic acid targets in clinical diagnostics, environmental monitoring, and food safety, to name a few examples [1-5]. However, these methods often require laborious and time-consuming steps like nucleic acid extraction and purification, which can lead to target loss, contamination, and increased assay time [6-8]. Traditional techniques like RT-PCR and RT-LAMP demand meticulous handling, hindering the development of rapid, sensitive, and affordable diagnostics [9, 10]. Rapid diagnostic tests, especially in outbreak situations, are crucial for timely disease management and public health interventions [11, 12]. Point-of-care (PoC) testing decentralizes diagnostics, bringing healthcare closer to patients in remote and underserved areas[13, 14]. Simplified sample preparation is key for developing portable and user-friendly diagnostic devices, which can improve patient outcomes and facilitate early intervention by eliminating the need for centralized labs and reducing turnaround times [12].

To overcome these limitations, there is a growing interest in developing nucleic acid amplification methods that can directly analyze complex samples without prior purification [15, 16]. Enzyme-free amplification approaches, such as toehold-mediated strand displacement reaction (TMSDR), offer a promising alternative to traditional enzymatic methods by allowing isothermal amplification, reducing the need for precise temperature control and specialized equipment [17-20]. By eliminating the use of enzymes, these methods can be more stable and less susceptible to variations under various environmental conditions. Reduced reliance on enzymes can lead to lower production costs and increased accessibility of diagnostic assays. Additionally, enzyme-free amplification can enhance the robustness and reproducibility of diagnostic tests. Kapadia et al. showcases the temperature resilient nature of TMSDR probes, which can be a significant advantage in PoC settings [20]. Moreover, TMSDR has demonstrated exceptional tolerance to common inhibitors present in various environmental and biological matrices. It remains effective in the presence of heparin used as anti-coagulant in blood and high concentrations of humic acid present in soil samples [21, 22]. Importantly, the robust performance of the TMSDR assay enables its direct application to bacterial lysates for 16S rRNA analysis. Given that the 16S rRNA gene is found in all bacteria species, targeting specific regions within this gene could be used for rapid and sensitive bacteria identification without the need for prior nucleic acid purification [23, 24].

This study explores a probe-based TMSDR amplification in the presence of various chaotropic agents, detergents, and lysis buffers. The assay was also performed in several biological and environmental samples to showcase the robustness and compatibility of the assay. A proof-of-concept assay targeting the 16S rRNA of *Escherichia coli* further confirmed specificity following tests against other bacteria with the probe.

## Materials and Methods

### Materials

All reagents used in this study were of analytical grade unless otherwise stated. Isothermal Amplification Buffer II was obtained from NEB. Guanidinium Hydrochloride (GDH) (Bioshop, Canada), Guanidine Isothiocyanate (GITC) (Bioshop, Canada), Sodium Dodecyl Sulfate (SDS) (Bioshop, Canada), and IGEPAL CA-630 (Sigma Aldrich, CAS: 9002-93-1) were procured from sources mentioned in parenthesis. The viral lysis buffer and environmental lysis buffer were obtained from Galenvs Science. Viral lysis buffer was from Viral RNA extraction kit (Galenvs Sciences) and environmental lysis buffer was obtained from Soil and Plant lysis kits (Galenvs Sciences). FAM and BHQ-1 modified oligonucleotide probes (Table S1) were synthesized at Galenvs using H16 synthesizers (K&A Labs) and purified by reverse phase high performance liquid chromatography (HPLC) to a purity of >85%. Two probes were designed, first one for the ORFa of SARS-CoV-2 variant B1.617.2 and second one targeting 16S rRNA of *E.coli* DH5α strain (Table S5.1). The probe designs were based on the parameters explained in this study [20]. Blood samples were obtained using PaxGene blood RNA collection tubes (Ref: 762165, Lot: 1131433), saliva from BioIVT (Cat: HUMANSALIVAUNN, Lot: HMN653021), and viral transport media (VTM) from Launchworks (Lot: 017377). Wastewater was collected from the Quebec City wastewater plant, while eggplant leaf and soil samples were obtained from Galenvs Sciences.

### Probe Preparation and TMSDR Assay

Probes were annealed using a thermal cycler to ensure optimal stem-loop formation and stem folding. in the assay. The probe was prepared using Isothermal Amplification Buffer II and then mixed with the various components as directed by the assay. The final concentration of the probe is 50 nM unless otherwise specified.

The TMSDR assay was conducted in a total volume of 50 µL in black, clear-bottom 96-well plates (Wuxi NEST Biotechnology Co., Ltd, Jiangsu, China). The reaction mixture consisted of Isothermal Amplification Buffer II, target nucleic acid, probe (50 nM unless otherwise specified), displacer (5 µM), and other components as required. All oligonucleotide stocks were prepared in water and the concentrations of the stocks were adjusted to 100 µM. Two target concentrations were assayed and the stock for these target nucleic acids were prepared in various buffer components and matrices in 10x concentrations and 5 µL of the prepared target was added into 40 µL (typically to obtain 5 nM or 1 nM final target concentration, as indicated in text) of prepared probe and 5 µL of 100x displacer. The target concentration in amplification wells is 5 nM and 1 nM unless otherwise stated. Reactions were incubated at room temperature (∼22°C) unless otherwise stated. Fluorescence kinetics data were collected at regular intervals using the microplate reader (Spectramax iD3, Molecular Devices or Tecan M1000 pro).

The TMSDR assay targeting 16S rRNA of *E.coli* differs from the assay explained above. The probe concentration for the assay was increased to 100 nM (final concentration) and 35 µL of bacterial lysate was introduced in the place of a target nucleic acid.

### Sample Preparation

Blood samples were diluted to a final concentration of 5% in the reaction mixture in viral lysis buffer. Saliva and Viral Transport Media (VTM) samples were also diluted in viral lysis buffer. Plant leaf samples (50 mg) were grinded using metal beads and resuspended with environmental lysis buffer. Soil samples (250 mg) were also resuspended with environmental lysis buffer. Wastewater sample was also diluted in environmental lysis buffer. All samples were spiked with target nucleic acid to achieve final concentrations of 5 nM and 1 nM. Here, DNA oligonucleotides mimicking ORF-1a gene of SARS-CoV-2 viral RNA (B.1.617.2 strain) was used as target nucleic acid.

Overnight cultures of three bacterial strains—*Escherichia coli* DH5α, *Methylobacterium extorquens*, and *Clavibacter michiganensis*—were used for cell lysate preparation. To break cells, for each strain, 900 µL of the culture was transferred into 2 mL screw-cap tubes pre-loaded with zirconium beads. The samples were then subjected to mechanical disruption using a tissue lyser for 40 seconds. This lysis step was repeated twice to ensure thorough cell disruption. Following the mechanical lysis, 100 µL of a 10% SDS solution was added to each tube to further facilitate cell breakage, enhance lysis efficiency and denature nucleases. The resulting lysate was used directly in the TMSDR assay without any additional processing.

### Data Analysis

All data analysis was performed using GraphPad Prism software. The assays were performed in three triplicates (i.e. a total of nine replicates per sample type) and for the significance of variance, two-way ANOVA was performed. The Standard Deviations (SD) were plotted as error bars.

## Results

### Enzyme-free amplification workflow

Unlike the traditional method, which requires time consuming purification and elution steps, the enzyme-free assay can be performed directly in lysis buffers (Figure 1). This streamlined workflow offers significant advantages, including reduced time-to-result and the potential for point-of-care applications. Moreover, by eliminating the need for nucleic acid purification, the enzyme-free method minimizes the handling of biohazardous materials, enhancing safety and convenience. In contrast, conventional methods like RT-PCR often rely on enzymatic amplification, which can be inhibited by detergents, alcohols, and chaotropic agents commonly found in lysis buffers. The enzyme-free nature of our assay overcomes these limitations, providing a more robust and versatile approach.

**Figure 1.**
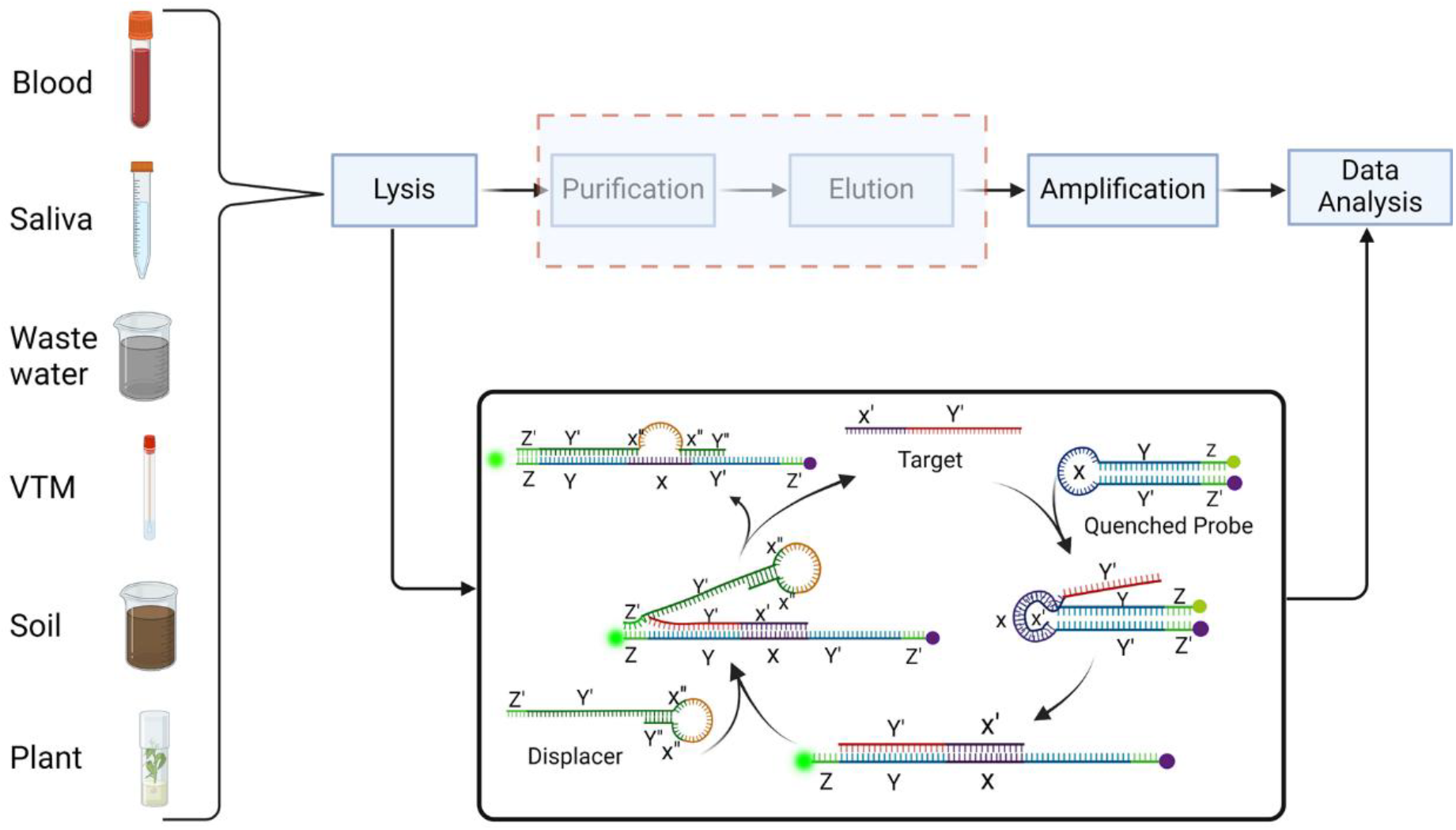
TMSDR workflow comparison with conventional RT-PCR workflow. The red dotted box highlights steps eliminated in the TMSDR workflow compared to RT-PCR.

### Enzyme free amplification in various lysis buffer and sample matrices

We performed a comprehensive analysis of the TMSDR assay’s performance under various conditions (Figure 2). In Figure 2 A, the effects of different buffer components and lysis buffers on amplification are compared. While all components had observable amplification, viral lysis buffer consistently exhibited superior performance, outperforming the standard control (i.e. assay in amplification buffer). Notably, 0.5 M GDH and 0.5 M GITC increased background, as evidenced by elevated fluorescence in the negative control, suggesting a lowered melting that allowed displacer based TMSDR in absence of target.

**Figure 2.**
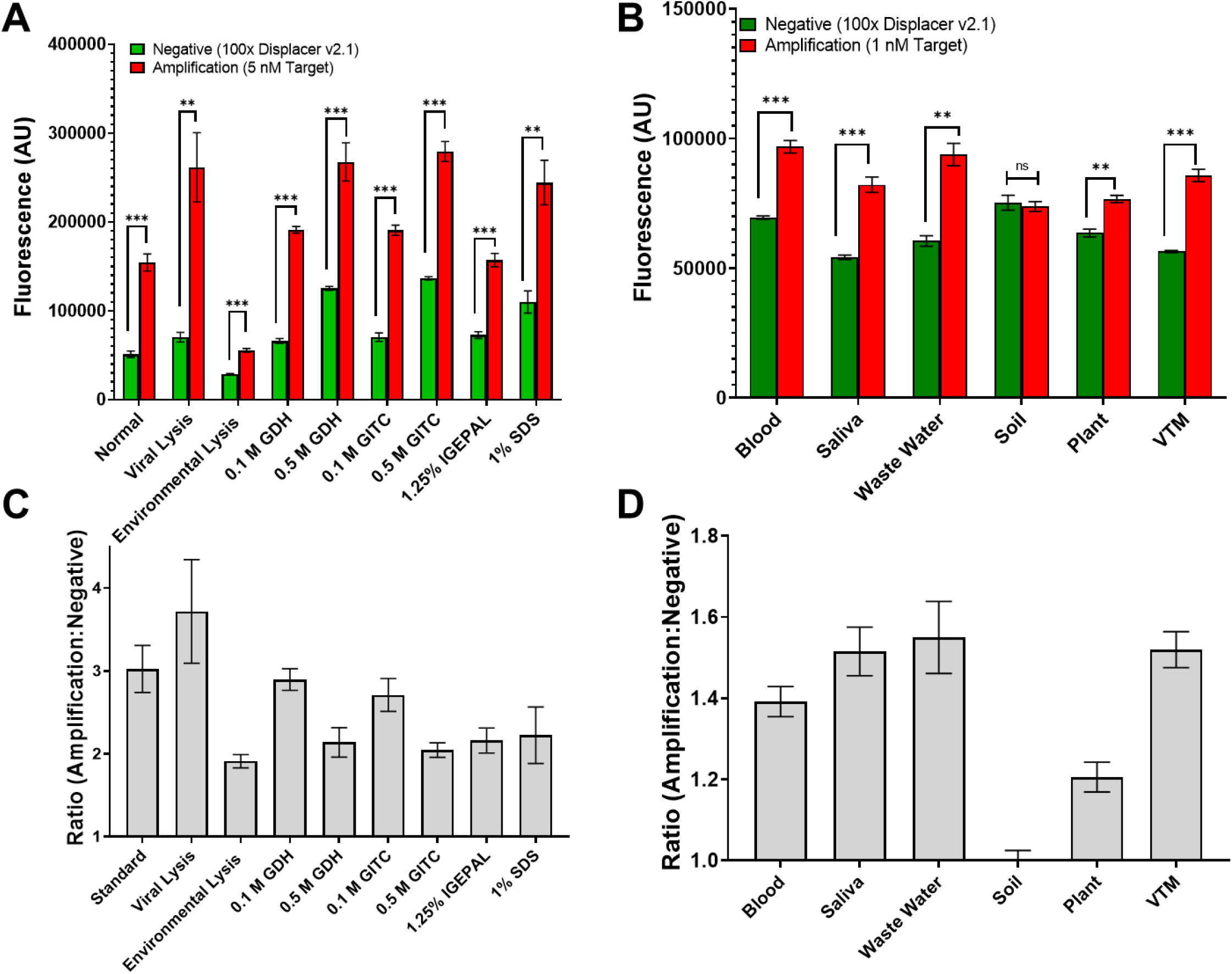
TMSDR amplification with various additives and in different matrices. The assay was conducted for 300 minutes, and fluorescence reads were taken every 2 minutes. **(A)** Comparative analysis of different buffers. Fluorescence levels of various conditions, each sample is represented by two bars: a green bar for the negative control (without target) and a red bar for the amplification (with 1 nM target strand). The assay was performed under standard conditions, and in the presence of viral lysis buffer, Environmental lysis buffer, 0.1 M GDH, 0.5 M GDH, 0.1 M GITC, 0.5 M GITC, 1.25 % IGEPAL, and 1% SDS. (P value: <0.05 =* ; <0.01=**; <0.001=***; Two-way ANOVA was performed on n=3) **(B)** Fluorescence levels across different sample matrices. Matrices tested: blood, saliva, wastewater, soil, plant, and VTM. Bars and significance as in (A). **(C** and **D)** Ratios comparing negative controls and amplification, calculated from Figure 1A and B. Each condition is represented by a single bar, allowing for a clear comparison of amplification efficiency across different additives and buffers **(C)** or sample matrices **(D)**.

Figure 2 B evaluates the assay’s sensitivity in different sample matrices using a target concentration of 1 nM. While most samples demonstrated significant amplification compared to negative controls, soil samples exhibited minimal fluorescence amplification, likely due to interference from the brown color and presence of humic acid [21]. The amplification data for a target concentration of 5 nM is provided in Supplementary Figure S1 A and B. To assess amplification efficiency more accurately, we determined the amplification-to-negative control ratio for each component and lysis buffer (Figure 2 C); and sample types (Figure 2 D). The results indicate that most components and lysis buffers exhibit favorable amplification ratios, except for Environmental sample buffer. As for sample types, most matrices showed significant amplification, except soil samples which displayed negligible ratios, suggesting interference from the matrix itself.

### Kinetic analysis of TMSDR assay in various matrices and with PCR inhibitors

We compared the effects of components and lysis buffers on k_obs_ and plateau (Figure 3). The standard and Viral lysis buffers exhibit the highest signal at the end of the reaction (after 5 hours), but high concentrations of chaotropic agents (0.5 M of GDH and GITC) increase the speed of TMSDR by several folds (Figure 3 A). While sample matrices have only a relatively minor impact on the final readout (except soil sample, Figure 2 B), with a similarly small impact on k_obs_, the reaction buffer components, particularly guanidine at 0.5 M, have the greatest impact on speed of reaction (Figure 3 B). Figure 3 B extends this analysis to various sample matrices, highlighting moderate impact of sample types. Saliva, VTM and wastewater exhibit the highest k_obs_ values, suggesting faster amplification kinetics. The Plateau values for these matrices also appear to be relatively high.

**Figure 3.**
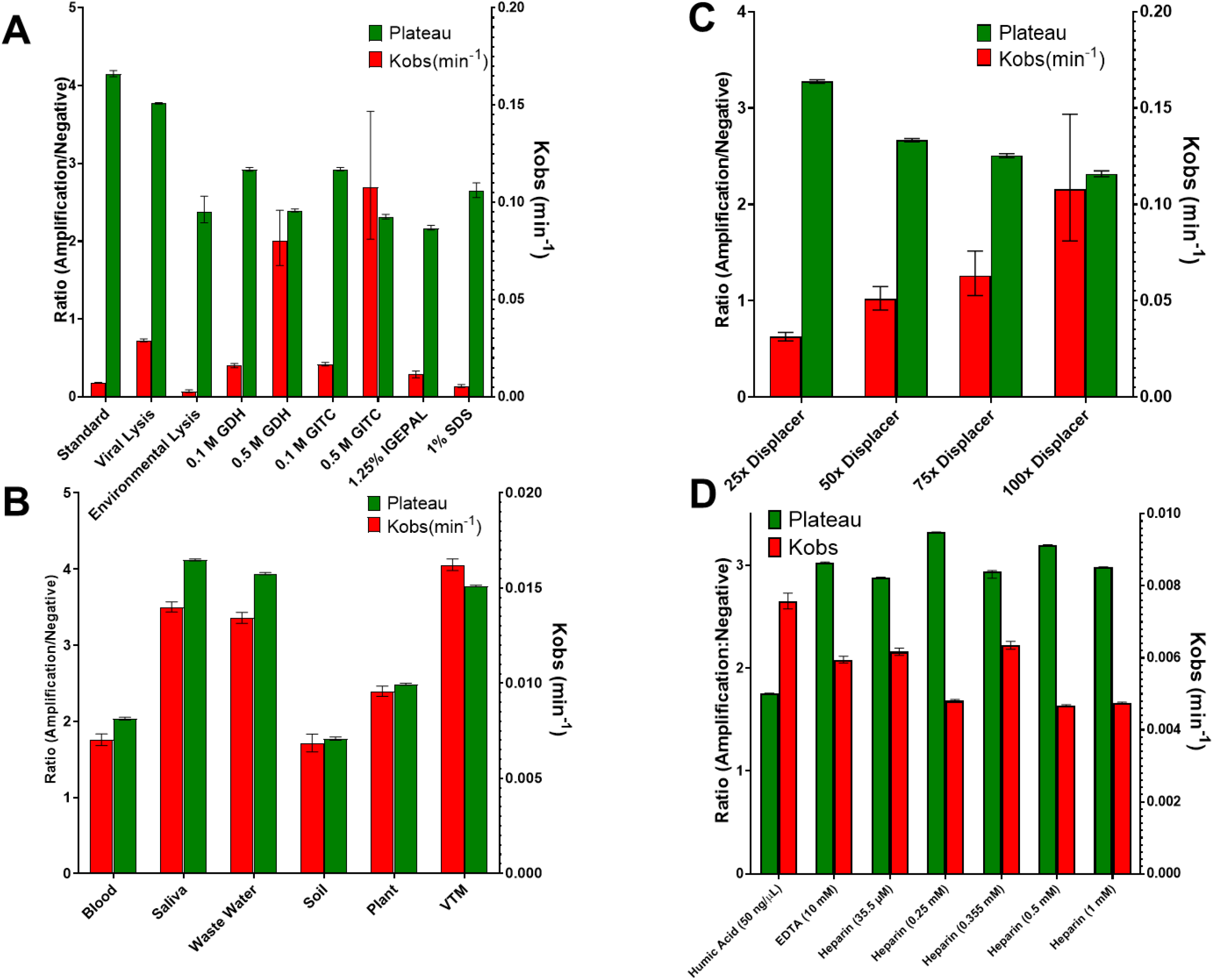
Amplification ratio and kinetic rate constants (k_obs_) for different components, lysis buffers, and sample matrices. **(A)** Comparison of amplification ratio and k_obs_ for various components and lysis buffers. **(B)** Comparison of amplification ratio and k_obs_ for different sample matrices. **(C)** Effect of displacer strand concentration on amplification ratio and k_obs_ (min^-1^) in the presence of GITC (0.5 M **(D)** Bar graph illustrates kobs and Plateau values of TMSDR in the presence of various inhibitors that are usually present in various biological matrices). Error bars indicate standard deviations from three independent experiments.

Increasing displacer concentration in the presence of 0.5 M GITC leads to higher k_obs_ values, indicating faster amplification rates. However, a moderate decrease in plateau values is observed at higher displacer concentrations (Figure 3 C), suggesting potential saturation effects and increased background in presence of too much displacer. Additionally, the variability in k_obs_ increases with higher displacer concentrations, aligning with the observations in Figure 3 A.

To further evaluate the robustness of our TMSDR assay in conditions known to be incompatible with polymerase-based amplification, we performed the assay in the presence notorious PCR inhibitors: heparin, humic acid and EDTA (Figure 3 D). TMSDR worked well in most assays without any large variations and no inhibitory effects were observed (Figure S3 A and B). However, humic acid adversely affected the assay and decreased plateau significantly. An oligonucleotide labelled with FAM at its 5′ end in presence of various concentrations of humic acids revealed that high concentrations of humic acid directly impact fluorescence (Figure S3 C), suggesting that TMSDR might actually occur normally in soil samples, but fluorescence is quenched.

Table 1 summarizes the data from the bar graphs (Figure 3 A) and a 5-star rating system is implemented to showcase the data. Chaotropic agents, GDH and GITC, at higher concentrations increase the rate of reaction for TMSDR, however it comes with higher background noise and larger variations. Viral lysis buffer showcases a good compromise as it has increased rate of reaction while maintaining high enough plateau value after 5 hours. While all samples show some degree of sensitivity, the viral lysis buffer has one of the highest signals (Plateau) with a k_obs_ higher than standard and excellent reproducibility, making it a preferred choice for TMSDR-based diagnostics. The star ratings and comments provide a quick summary of the performance of each sample, helping identify the best and worst performers effectively.

**Table 1.**
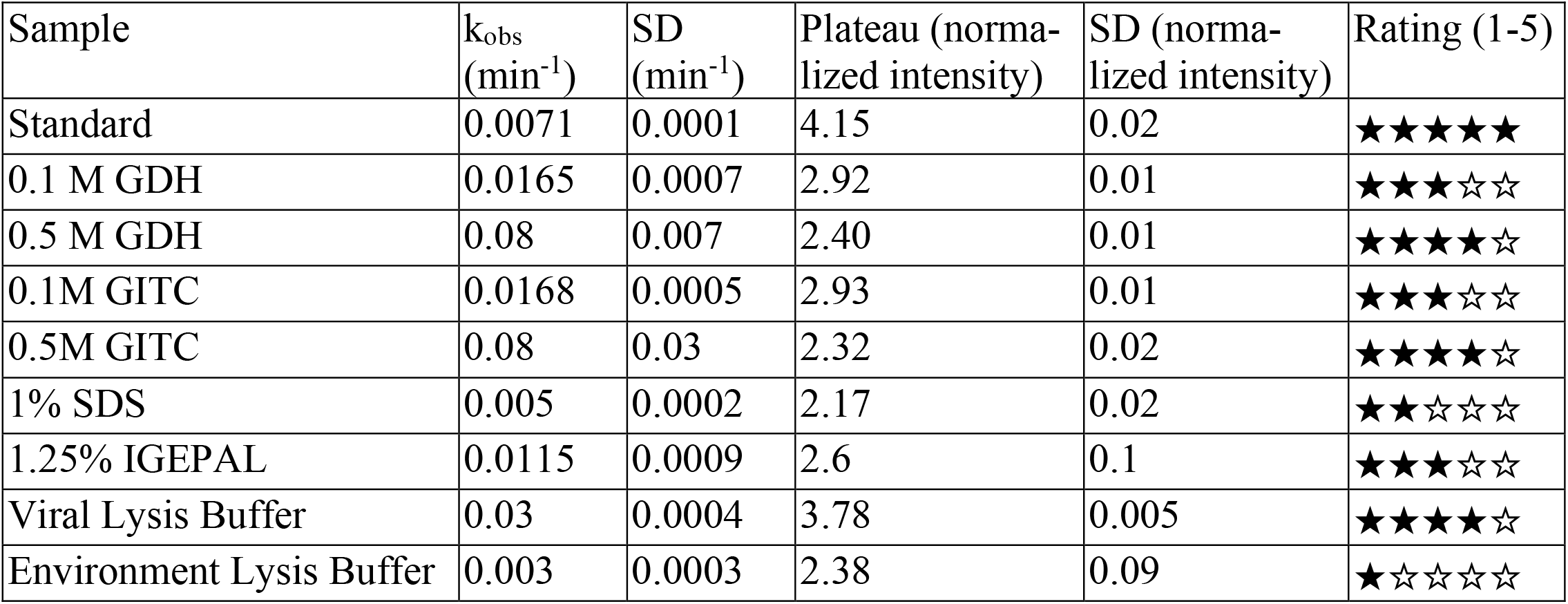
Summary of TMSDR with various buffers. All values were derived from curves generated over a 300-minute period, with readings taken every 2 minutes. The standard deviations (SD) for k_obs_ and plateau values are included to indicate the variability and reliability of the measurements. Star ratings summarize the performance, with higher ratings indicating overall best compromises between kinetics and robustness.

Table 2 summarizes the data derived from Figure 3 B and D, utilizing a similar 5-star rating system to highlight amplification trends across various sample types. The TMSDR amplification method is compatible with a wide range of biological and environmental sample types as well as inhibitors present in various sample types. However, the color of the sample can impact the fluorescence readout, as evidenced by the soil sample and effect of humic acid on fluorescence. Wastewater and plant samples showed good sensitivity and stability, with k_obs_ of 0.013 min^-1^ and 0.0096 min^-1^, respectively, and stable plateau values of 3.96 and 2.49. These results suggest that TMSDR can reliably detect target analytes in these complex matrices, making it a versatile method for environmental monitoring and agricultural applications. In contrast, soil samples presented the lowest sensitivity, with a k_obs_ of 0.0068 min^-1^ and a plateau of 1.77. The higher standard deviations observed in soil samples indicate greater variability, reflecting the complex nature of soil matrices and the challenges of detecting fluorescence in samples with humic acid.

**Table 2.**
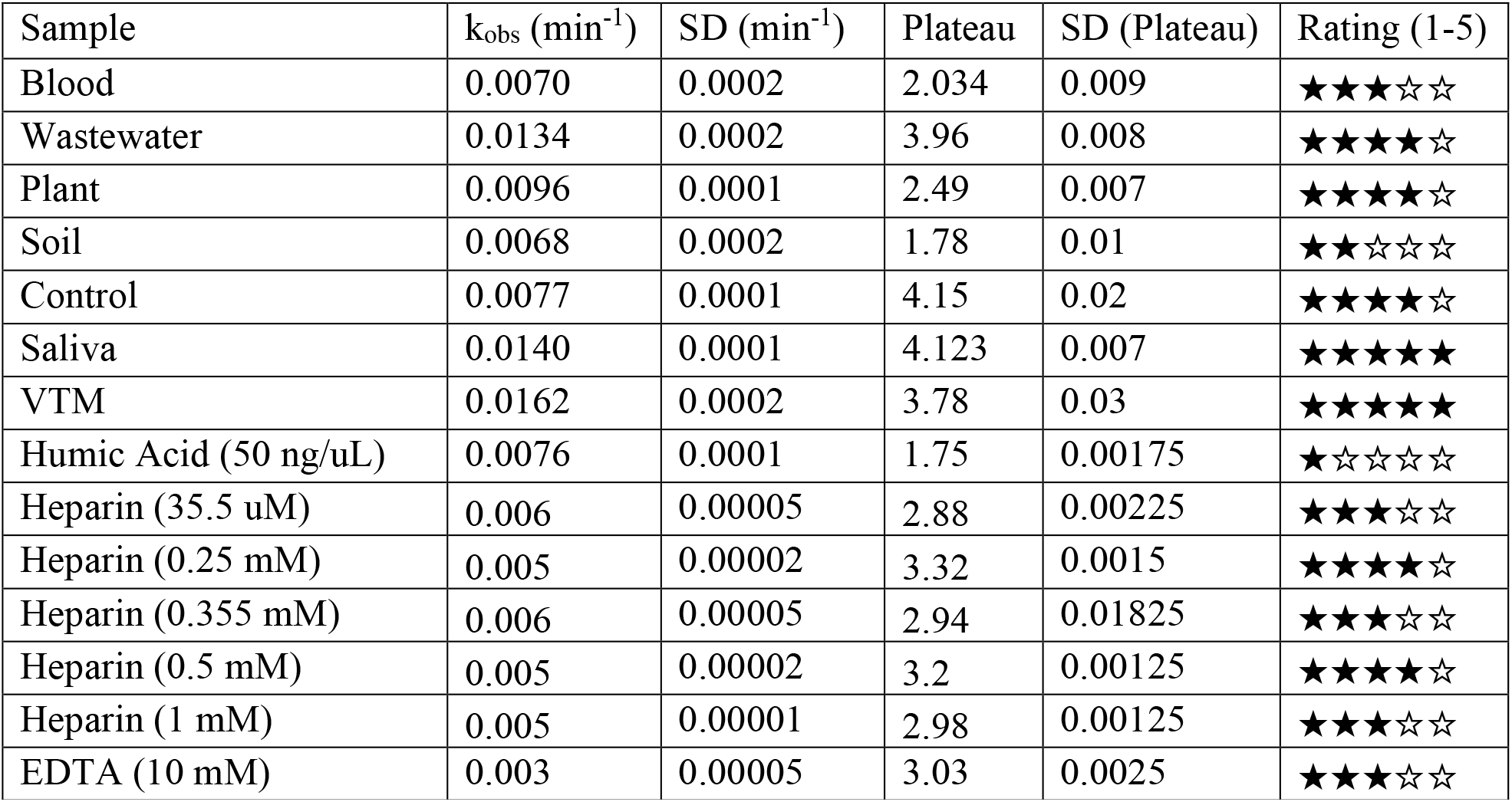
Summary of TMSDR in various biological and environmental samples. All values were derived from curves generated over a 300-minute period, with readings taken every 2 minutes. The standard deviations (SD) for k_obs_ and plateau values are included to indicate the variability and reliability of the measurements. Star ratings summarize which sample types are easier to assay.

Blood samples displayed moderate performance, with a k_obs_ of 0.007 min^-1^ and a plateau of 2.034, indicating adequate sensitivity but less robustness compared to top-performing samples. Heparin, and EDTA, common anti-coagulants used in blood samples [22, 25], do not have any adverse effect on TMSDR. Saliva samples also exhibited excellent performance, closely matching VTM sample with consistent results and small standard deviations.

Overall, the star ratings assigned to each sample type provide a quick visual summary of our TMSDR assay performance with each, with VTM and saliva earning top ratings for their excellent sensitivity and robustness. TMSDR assay performance was also good with wastewater and plant samples, whereas blood and soil samples exhibited moderate to lower performance. The calculated standard deviations further highlight the reliability of the measurements, with smaller deviations indicating more consistent results. This comprehensive analysis underscores the potential of TMSDR-based biosensors as a versatile and reliable diagnostic tool across various sample types, particularly for applications requiring rapid and sensitive detection without extensive sample processing.

### TMSDR assay targeting 16S rRNA of *E.coli*

TMSDR assays were performed using lysates from three bacterial strains—*E. coli, M. extorquens*, and *C. michiganensis*—as described in the Materials and Methods section. A probe and displacer sequence targeting the 16S rRNA of *E.coli* was engineered based on the parameters described in [20]. The target sequence of 16S rRNA is illustrated in Figure 4 A. *M. extorquens* and *C. michiganensis* target sequences are also showcased, and the mismatch nucleotides were highlighted in red (Figure 4 A). Robust amplification was observed exclusively with the *E. coli* lysate, whereas lysates from *M. extorquens* and *C. michiganensis* produced signals that were comparable to the negative control (Figure 4 B). As shown in Figure 4 C, end-point fluorescence measurements confirm the selective amplification in *E. coli*. Furthermore, Figure 4 D presents the ratio of the amplification signal to the negative control, reinforcing the enhanced performance of the assay in *E. coli*, while Figure 4 E illustrates the corresponding end-point ratio. Collectively, these results demonstrate the specificity of the TMSDR assay for bacterial targets under the conditions tested.

**Figure 4.**
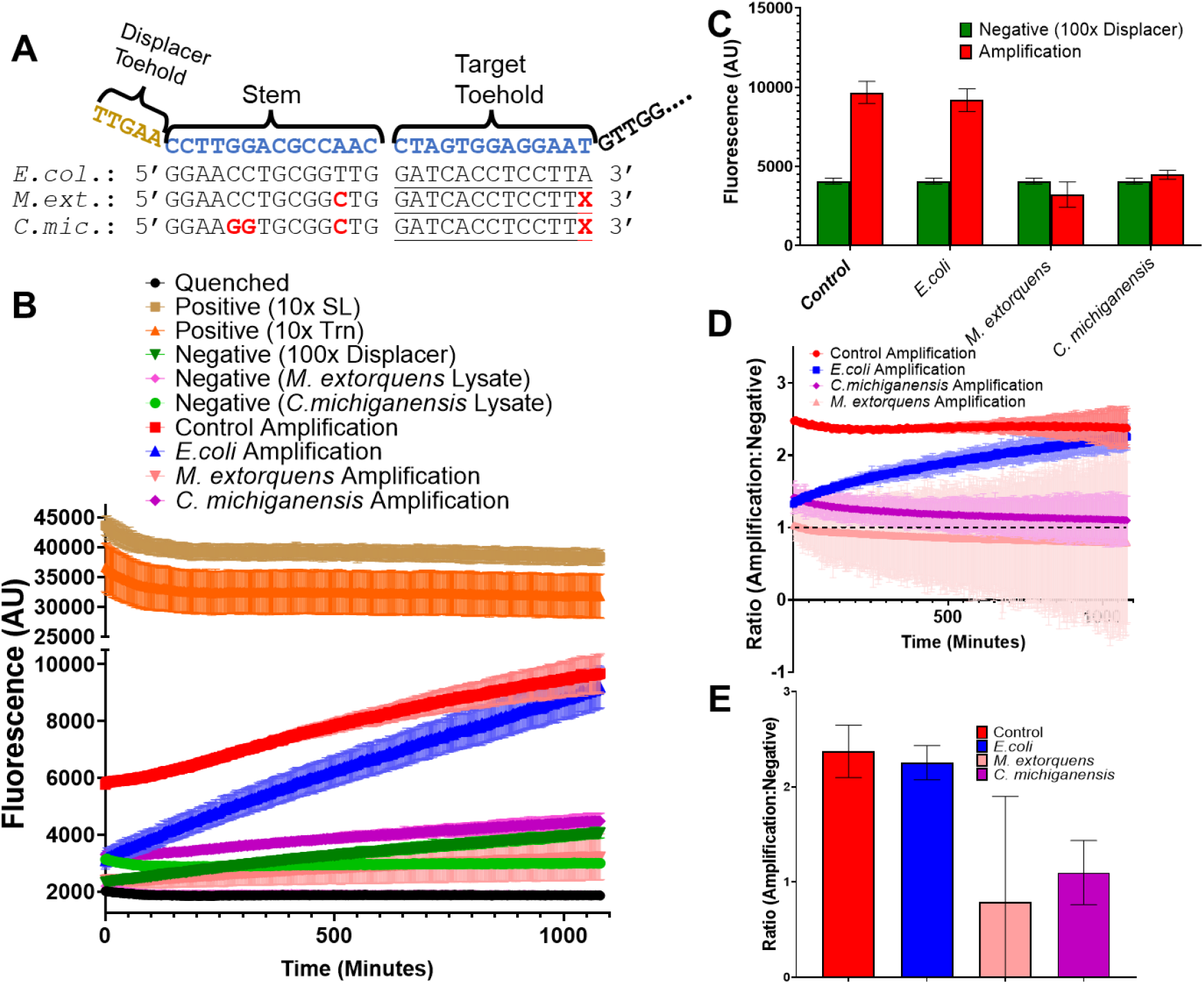
TMSDR assay using a probe targeting the 16S rRNA of *E. coli* compared with two other bacterial lysates. **(A)** Probe design and target sequence of *E.coli., M. extorquens* and *C. michiganensis*. Red nucleotides illustrate mutations in the target sequence. Target binding probe sequence is illustrated in blue fonts. **(B)** Graph illustrates TMSDR assay in the presence of various bacterial lysate **(C)** Comparison of end point raw fluorescence data extracted from **(B)** with negative control. **(D)** TMSDR assay ratio curve comparison of *E.coli, M. extorquens* and *C. michiganensis*. Control amplification was used as the benchmark. **(E)** Comparison of end point amplification ratio of control, *E.coli, M. extorquens* and *C. michiganensis*.

## Discussion

The comparative scheme presented in Figure 1 highlights the advantages of the TMSDR workflow over the conventional RT-PCR workflow [26]. By skipping several labor-intensive steps, as indicated by the grayed-out area in the red dotted box, TMSDR offers a streamlined and efficient method for nucleic acid amplification [27]. This reduces the time and resources required for sample processing and minimizes the risk of contamination [27]. The use of a fluorescence plate reader for detecting the amplified signal further enhances the sensitivity and accuracy of the assay.

By eliminating the need for nucleic acid purification, the TMSDR assay reduces sample handling time and minimizes the risk of sample degradation [27]. Additionally, the enzyme-free nature of the TMSDR assay makes it less susceptible to inhibition by contaminants commonly found in biological as well as environmental samples [28] and increases the shelf-life of the assay in point of care setting [29]. TMSDR effectively amplifies the target nucleic acid across a range of additives and buffers, including viral lysis buffer, soil lysis buffer, GDH, GITC, IGEPAL, and SDS. The assay retains functionality even in the presence of heparin and EDTA, commonly used in blood samples as anti-coagulant [22, 25]. This adaptability is crucial for applications in diverse fields, from clinical diagnostics to environmental monitoring [30].

However, matrix effects, such as the interference from the presence of humic acid in soil samples, can influence fluorescence signal detection (Figure 2 and 3 and Supp. Materials Figure S3 C). While soil samples were the ones where sensitivity was the lowest, due to the direct impact of soil material on fluorescent readout, amplification was still possible in spite of the notorious presence of PCR inhibitors in soil [21]. This could be further alleviated, for instance by changing the detection platform from fluorescence to electrochemistry.

The evaluation of amplification ratio and k_obs_ provides deeper insights into the performance of TMSDR. We assayed the displacer concentration in the presence of chaotropic agent GITC (Figure 3 C). While higher displacer concentrations can enhance amplification kinetics in the presence of 0.5 M GITC, excessive concentrations may lead to saturation effects and reduced plateau values (Figure 3 C and Supp. Materials Figure S2). From these results, displacer concentrations need to be adjusted according to the concentrations of chaotropic agents, as both have significant impact on amplification efficiency and sensitivity.

Chaotropic agents (GDH, GITC and viral lysis buffer) clearly increased the rate for signal occurrence compared to detergents (IGEPAL, SDS), but had little impact on the end-point signal. However, the important contribution is not as much the slight improvement in assay results, rather than the fact that the assay is compatible with these commonly used denaturing buffers and chemicals that are otherwise completely incompatible with enzyme-based amplification methods [30]. This was further extended to biological and environmental matrices. Similar results were obtained for Saliva, VTM and waste-water samples. Plant and soil samples partially interfered with fluorescence readout of the assay.

The TMSDR assay, employing a probe-displacer pair specifically designed to target the 16S rRNA of *Escherichia coli*, clearly demonstrates its applicability on real biological samples. When applied to lysates from *Methylorubrum extorquens* and *Clavibacter michiganensis*, the assay produced only background-level amplification, underscoring its high specificity. Notably, the 16S rRNA sequences of *E. coli* and *M. extorquens* differ by only a single nucleotide, further attesting to the precision of the TMSDR approach (Figure 4). Given that 16S rRNA is widely used to identify bacteria, this further validates the precision of the TMSDR approach [24]. Moreover, the TMSDR assay’s ability to tolerate up to 1% SDS—a concentration ten times higher than the tolerance limit of the ionic detergent-tolerant assay reported by Nakanish et al., 2022, and significantly greater than the 0.001–0.003% SDS tolerated by most conventional PCRs—sets it apart as a robust solution for direct analysis of bacterial lysates without any purification [31]. While this study was designed to detect the 16S rRNA of *E.coli.*, the design parameters can be applied to any nucleic acid sequence. Despite its many advantages, this study also highlights several limitations that warrant further investigation and optimization. A key drawback of the current TMSDR setup is its relatively low sensitivity; it can detect target nucleic acids only at final concentrations of approximately 1 nM, which is considerably less sensitive than conventional enzymatic amplification methods [32, 33]. We managed to detect bacterial RNA without amplification, but rRNA is very abundant and we used much more bacteria than what is used in typical diagnostic tests, underscoring this sensitivity gap [34]. Additionally, while the assay is robust against common PCR inhibitors such as polyphenols present in plants and heme present in blood, interference from humic acid in soil samples remains problematic, as it appears to directly affect fluorescence readout (Supp. Materials Figure S3 C) [22, 35]. This limitation could potentially be addressed by switching to an electrochemical detection platform or by employing alternative fluorophores that are less susceptible to humic acid interference [36]. Further improvements in sensitivity might also be achieved through absorbance-based assays or by functionalizing probes with gold nanoparticles to enable aggregation-based detection [37, 38]. These challenges point to areas for future refinement, aiming to enhance the TMSDR assay’s performance for broader diagnostic applications. Overall, the results demonstrate the potential of TMSDR as a versatile and reliable diagnostic tool, particularly in applications requiring quick detection without extensive sample processing.

## Conclusion

In conclusion, the results of this study confirm that the TMSDR method—coupled with an optimized probe-displacer system—provides a robust, adaptable, and efficient platform for nucleic acid detection. The enzyme-free nature of TMSDR and its streamlined workflow dramatically reduce the need for complex sample preparation, thereby minimizing handling time, target loss, and the risk of contamination. The TMSDR assay demonstrates remarkable robustness, tolerating up to 1% SDS and functioning across various reagents, including lysis buffers, chaotropic agents, and detergents. In contrast, conventional PCR methods only tolerate 0.001–0.003% SDS while even ionic detergent-tolerant assays reach only 0.1% SDS, highlighting the superior performance of TMSDR.This tolerance enables direct analysis of complex samples, such as bacterial lysates, without any purification. Moreover, the assay maintains high performance in the face of common PCR inhibitors found in biological and environmental samples—such as polyphenols, humic acid, heparin, and EDTA—further broadening its applicability. The selective amplification of the 16S rRNA target from *Escherichia coli* contrasted with background signals from other bacterial lysates and attests to the high specificity of TMSDR; even a single nucleotide difference between the 16S rRNA sequences of *E. coli* and *Methylorubrum extorquens* is discernible, highlighting its potential for detecting single nucleotide polymorphisms. Although this work has focused on a 16S rRNA target, the modular design of the TMSDR assay can be readily adapted for any nucleic acid target, positioning it as a promising tool for future development of rapid, on-site diagnostics and a host of applications in both clinical and environmental settings. Future research should further enhance detection limits, ultimately broadening the utility of this innovative diagnostic platform.

## Author’s contributions

J.B.K, J.D and J.P designed all the experiments and took part in writing and editing the manuscript.

K.D.S.N prepared bacteria samples and did bacteria detection assay. J.B.K. performed all the other experiments.

## Conflict of interest

Galenvs Sciences is an oligonucleotide provider and some of the materials and designs provided by Galenvs sciences are part of a provisional patent filed by Galenvs Sciences.

## Notes

### Competing Interest Statement

The authors have declared no competing interest.

### Funding

- MITACS ACCELERATED GRANT

