## Supplementary material for "A purification-free nucleic acid amplification platform for diverse samples": Supp. Data

**Supplementary material for TMSDR: A purification free amplification platform for diverse samples**

1: INRS, Centre Armand-Frappier Santé Biotechnologie, 531 boul. Des prairies, Laval, QC, H7V 1B7

2: Galenvs Sciences: 6750 Rue Hutchison suit 201, Montreal, QC, H3N 1Y4

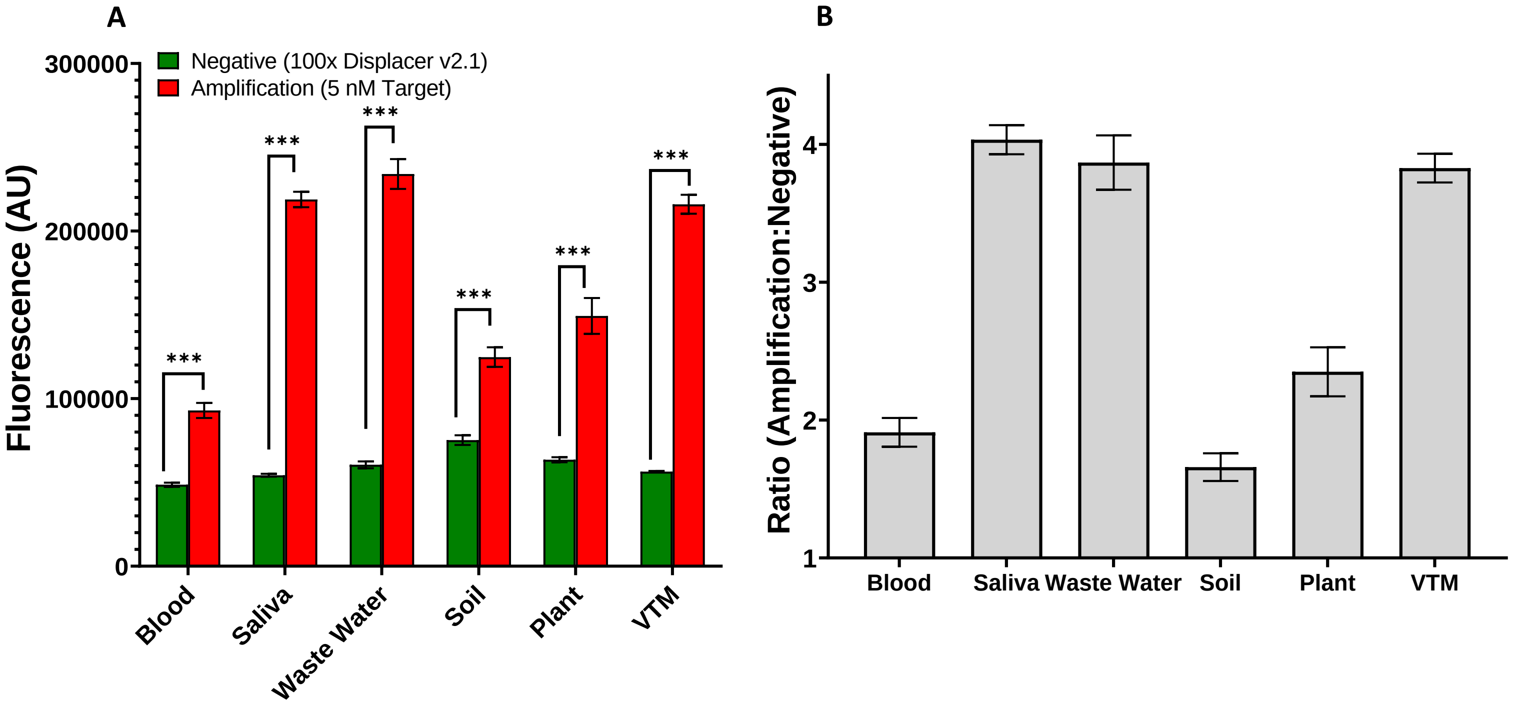


**Figure S1.** Comparison of amplification in various environmental and biological samples with 5 nM target. All the experiments were performed in independent triplicates. **(A)** This bar graph displays raw fluorescence data, where red bars represent amplification (probe + displacer + target) and green bars indicate the negative control (probe + displacer, without target). The graph provides a clear comparison of fluorescence levels across different samples. (P value: <0.05 =* ; <0.01=**;  <0.001=***; Two-way ANOVA performed with n=3 replicates).  **(B)** This bar graph illustrates the ratio of amplification to the negative control across different samples, providing a precise comparison of amplification levels.

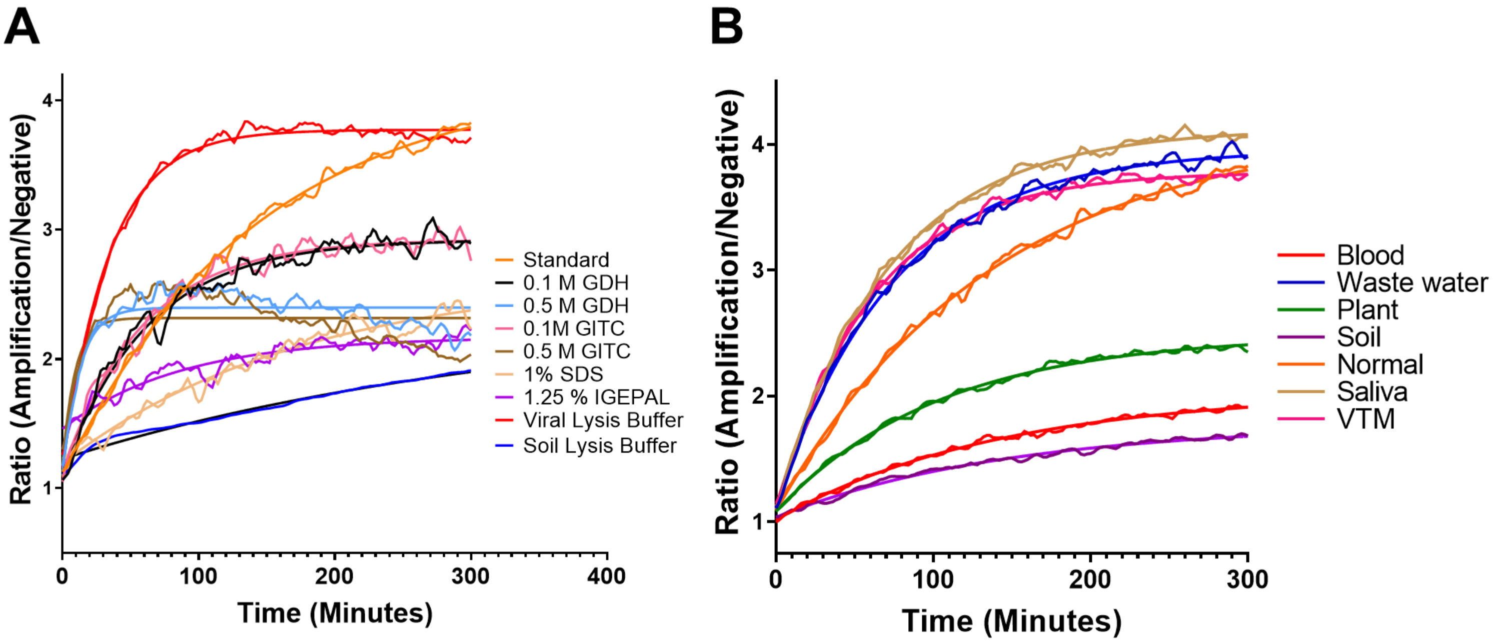


**Figure S2.** Kinetic data of TMSDR amplification in various buffers additives and matrices. The curves were smoothened, and the assay was conducted for 300 minutes with readings taken every 2 minutes at room temperature. **(A)** TMSDR assay in the presence of various lysis buffers, chaotropic agents and detergents. **(B)** TMSDR assay with environmental and biological matrices.

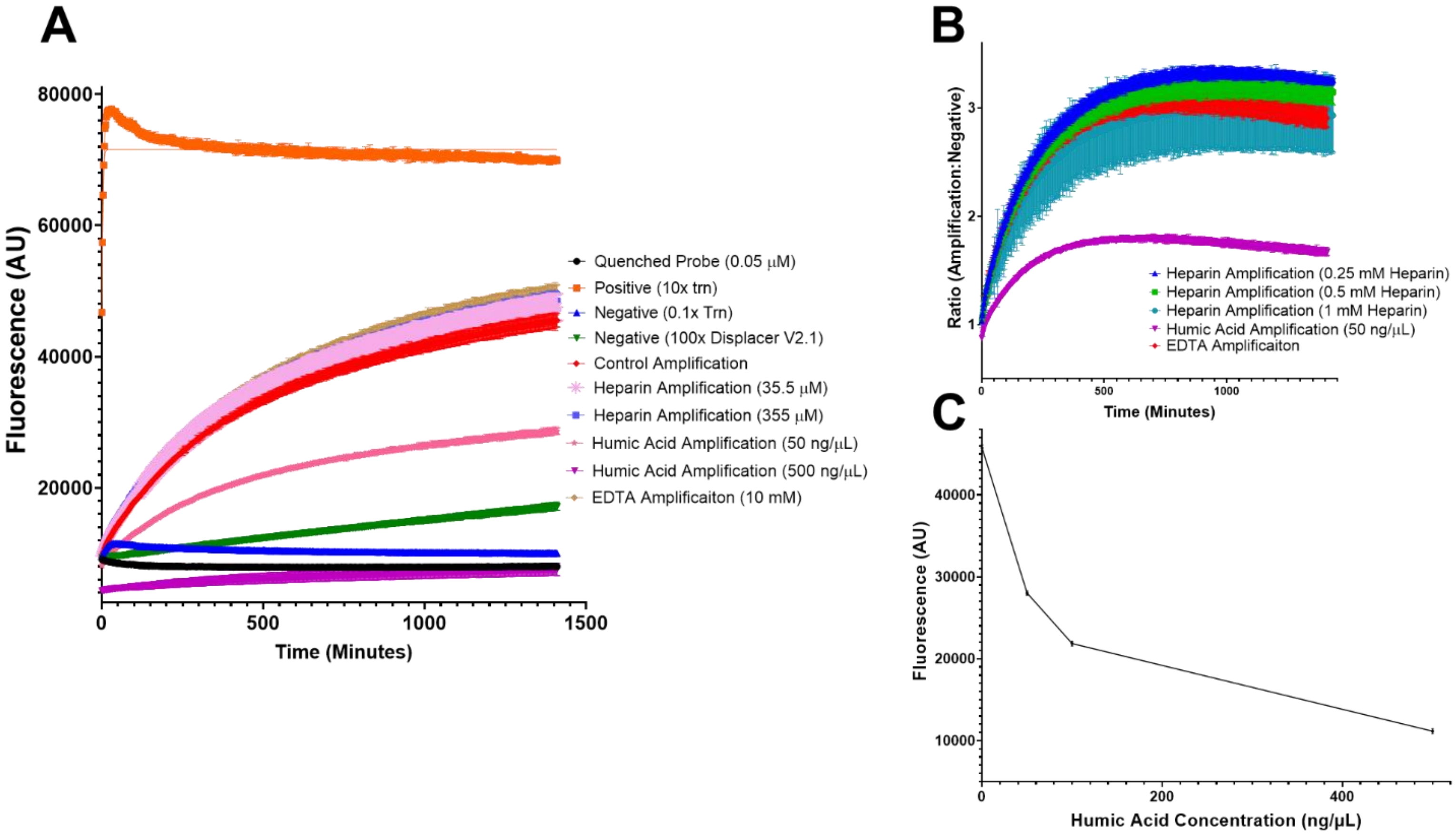


**Figure S3.** TMSDR assay in the presence of various PCR inhibitors and effect of humic acid on fluorescence. **(A)** TMSDR assay with various concentrations of heparin, humic acid and EDTA. **(B)** Ratio of amplification to negative control calculated from (A) for realistic comparison of TMSDR amplification **(C)** Effect of humic acid concentration on the oligonucleotide modified with FAM at 5’ end. Various concentrations of humic acids were utilized. Note that all tested concentrations of inhibitors were compatible with a clear amplification aside from concentrations of humic acid > 100 ng/μL.

| Sequence Name | Sequence (5′→ 3′) |
|---|---|
| Probe_SARS-CoV-2 | ACGTGGCTTTGGAGACTCCGTGGAGCTCTGATAAGACCTCCTCCACGGAGTCTCCAAAGCCACGT |
| Displacer_SARS-CoV-2 | CCGTGGAGGAGAAAAAAAAAGAGCTCCACGGAGTCTCCAAAGCCACGT |
| Trn_Target_SARS-CoV-2 | GAGGTCTTATCAGAGCTCCACGGAGTCTCCAAAGC |
| SARS-CoV-2 Target | GAGGTCTTATCAGAGCTCCACGGAGTCTCCAAAGC |
| Probe_16S rRNA | AACTTGGAACCTGCGGTTGTAAGGAGGTGATCCAACCGCAGGTTCCAAGTT |
| Displacer_16S rRNA | AACTTGGAACCTGCGGTTGGATAAAAAAATAGCAAC |
| Target_16S rRNA | GGAACCUGCGGUUGGAUCACCUCCUUA |

**Table S1.** Sequences used in the experiments. Probe_SARS-CoV-2 and Probe_16S rRNA sequences are modified with fluorophore (FAM) at 5′ end and quencher (BHQ-1) at 3′ end.
